# Menopause status- and sex-related differences in age associations with spatial context memory and white matter microstructure at midlife

**DOI:** 10.1101/2024.04.24.589653

**Authors:** Rikki Lissaman, Sricharana Rajagopal, Julia Kearley, Stamatoula Pasvanis, M. Natasha Rajah

## Abstract

Decline in spatial context memory emerges in midlife, the time when most females transition from pre-to post-menopause. Recent evidence suggests that, among post-menopausal females, advanced age is associated with functional brain alterations and lower spatial context memory. However, it is unknown whether similar effects are evident for white matter (WM) and, moreover, whether such effects contribute to sex differences at midlife. To address this, we conducted a study on 96 cognitively unimpaired middle-aged adults (30 males, 32 pre-menopausal females, 34 post-menopausal females). Spatial context memory was assessed using a face-location memory paradigm, while WM microstructure was assessed using diffusion tensor imaging. Behaviorally, advanced age was associated with lower spatial context memory in post-menopausal females but not pre-menopausal females or males. Additionally, advanced age was associated with microstructural variability in predominantly frontal WM (e.g., anterior corona radiata, genu of corpus callosum), which was related to lower spatial context memory among post-menopausal females. Our findings suggest that post-menopausal status enhances vulnerability to age effects on the brain’s WM and episodic memory.

## 1 Introduction

Aging is associated with decline in episodic memory – our ability to encode and retrieve past events in rich contextual detail (Grady, 2012; Tulving, 1972). Accumulating evidence indicates that age-related episodic memory decline, assessed using item-location spatial context memory tasks, begins in midlife and is associated with functional alterations in prefrontal and occipito-temporal cortical regions (Ankudowich et al., 2016; Cansino, 2009; Cansino et al., 2012; Kwon et al., 2016). Decline in spatial context memory is also linked to smaller posterior hippocampal volumes (Snytte et al., 2022) and greater functional connectivity between posterior hippocampus and fronto-parietal/occipital regions (Ankudowich et al., 2019), the latter of which seems to emerge at midlife. Collectively, these findings highlight midlife as a critical period in which episodic memory decline and associated structural/functional brain changes first arise and underscore the sensitivity of spatial context memory tasks in detecting this decline.

Notably, midlife is the time at which most females experience spontaneous menopause and transition from pre-to post-menopause (Harlow et al., 2012). Many females report cognitive difficulties, including memory problems, during the menopausal transition (Greendale et al., 2020). This raises the possibility that prior reports of age-related spatial context memory decline at midlife may be driven (in part) by post-menopausal females, and that menopause status may contribute to the presence of sex differences in the effect of age on episodic memory at this critical period. Indeed, recent studies have observed sex differences in the relationship between chronological age and spatial context memory-related functional activity and connectivity (Subramaniapillai et al., 2019, 2022; Wang et al., 2022).

To our knowledge, only two neuroimaging studies have explored differences in the neural correlates of episodic memory as a function of menopause status, both using fMRI (Crestol et al., 2023; Jacobs et al., 2016). In the first such study, Jacobs et al. (2016) found that post-menopausal females exhibited lower left hippocampal activity and greater bilateral hippocampal connectivity during verbal encoding than pre-menopausal females, although no behavioral effects were observed on subsequent retrieval. Interestingly, among post-menopausal females specifically, Jacobs et al. further reported that “high” performers on an out-of-scanner associative memory task demonstrated the lowest levels of bilateral hippocampal connectivity, suggesting that menopause-related increases in functional connectivity are linked to episodic memory disruption. In this study, however, menopausal groups were age-matched, an approach which assumes chronological and reproductive aging to be independent processes. By contrast, Crestol et al. recently examined associations between age and spatial context memory-related functional activity within pre- and post-menopausal females, respectively. The authors observed that, in post-menopausal females only, advanced age was associated with lower activation in occipito-temporal and parahippocampal cortices during encoding and retrieval, which in turn was associated with lower spatial context memory. This finding indicates that menopause may act as an inflection point in some females, after which vulnerability to age effects on the brain and cognition increase. In this view, chronological and reproductive aging constitute synergistic, rather than independent, processes. However, as Crestol et al.’s pre- and post-menopausal groups differed in age, it is also possible that their results reflected more general early-vs. late-midlife age effects, which may likewise be evident in males.

Neuroimaging evidence suggests that sex- and menopause-related differences are evident in the micro-structural properties of the brain’s white matter (WM), as well as its association with chronological age. The brain’s WM is primarily comprised of myelinated axons bundles, or tracts, supporting the efficient transmission of information between regions. Using diffusion tensor imaging (DTI), it is possible to probe the microstructural properties of WM tracts in vivo (Soares et al., 2013). The two most common DTI measures are fractional anisotropy (FA), which represents the degree to which diffusion is constrained in a given direction, and mean diffusivity (MD), which represents the overall diffusion rate. While several factors (e.g., myelination, axon density, axon diameter) can affect these measures (Jones et al., 2013), low FA and high MD are often used as indices of poorer microstructural health or “integrity”. Several large-scale studies have reported age and sex effects on WM throughout the brain, with interactions occasionally evident (e.g., Ritchie et al., 2018; Lawrence et al., 2021). For example, Isaac Tseng et al. (2021) found that age negatively impacted the microstructural properties of more tracts among females than males, which the authors hypothesized may be linked to menopause. Relatedly, one recent study observed lower FA in the external capsule among post-compared to peri-menopausal females, with further differences evident between pre-, peri-, post-menopausal females and age-matched males (Mosconi et al., 2021). However, since this study did not examine age effects within groups or include cognitive measures, it is unclear whether the relationship between age and WM microstructure differs based on menopause status and whether this has any implications for episodic memory at midlife.

In this study, we investigate the association between age and spatial context memory in pre- and post-menopausal females. We utilize data from a face-location spatial context memory paradigm that is sensitive to chronological age effects at midlife, particularly among post-menopausal females (Crestol et al., 2023). We thus hypothesize that age will be negatively associated with spatial context memory among post-but not pre-menopausal females. Moreover, as we theorize that this association is influenced by pre-/post-menopause effects and not more general early-/late-midlife effects, we hypothesize that age will be negatively associated with spatial context memory in females but not males. Given prior neuroimaging evidence, we further hypothesize that there will be menopause status- and sex-related differences in the relationship between age and DTI-derived measures of WM microstructure and, in addition, that these differences will contribute to individual differences in spatial context memory.

Methods

## 2 Materials and Methods

### 2.1 Participants

One-hundred and seventeen cognitively unimpaired middle-aged adults (39.55-65.46 years, *M* = 52.02, *SD* = 6.84) participated in this study. Of these, 34 were males (*M*_*age*_ = 53.70, *SD*_*age*_ = 5.65, age range: 43.84-65.46; *M*_*EDU*_ = 16.54; Ethnicity: 2 Arab, 1 Black, 1 Chinese, 1 Latin American, 29 White), 37 were pre-menopausal females (*M*_*age*_ = 44.38, *SD*_*age*_ = 3.03, age range: 39.55-53.30; *M*_*EDU*_ = 16.34; Ethnicity: 1 Black, 1 Chinese, 1 Latin American, 1 South Asian, 32 White, 1 White & Latin American), and 46 were post-menopausal females (*M*_*age*_ = 56.92, *SD*_*age*_ = 3.89, age range: 47.26-65.14; *M*_*EDU*_ = 15.16; Ethnicity: 1 Arab, 1 Black, 1 Chinese, 1 Indigenous, 1 North African, 1 South Asian, 40 White). Pre-/post-menopause status was based on the Stages of Reproductive Aging Workshop + 10 (STRAW+10) criteria (Harlow et al., 2012), described below (Section 2.3). All participants signed informed consent forms and were compensated for their time. The research ethics board of the Montreal West Island Integrated University Health and Social Services Centre (subcommittee for mental health and neuroscience) approved the study procedures.

### 2.2 Procedure

Participants were recruited via online advertisements and flyers posted around the Montreal area. At enrollment, all participants signed an online consent form and completed a series of questionnaires. The information provided was used to screen participants for the larger, ongoing Brain Health at Midlife and Menopause (BHAMM) study, of which the current study is a sub-study (Crestol et al., 2023). To be included, participants had to possess a high school diploma, agree to provide blood samples for endocrine assessment to validate self-reported menopause status, and be in good general health. Exclusion criteria were: current use of hormone replacement therapy; bilateral oophorectomy; untreated cataracts/glaucoma/age-related maculopathy; uncontrolled hypertension; untreated high cholesterol; diabetes; history of estrogen-related cancers; chemotherapy; neurological disease or history of serious head injury; history of major psychiatric disorders; claustrophobia; history of substance abuse disorder; currently smoking more than 40 cigarettes per day; did not meet MRI safety requirements. For this sub-study, we excluded females who were pregnant, peri-menopausal, or whose menopause status was indeterminate (based on self-report and hormones). Post-menopausal females were also excluded if they were using hormonal birth control, while pre-menopausal females were excluded only if they were using hormonal birth control for reasons other than contraception. We additionally excluded participants if their BMI was greater than 40 or if their chronological age was more than 2.5 standard deviations above/below their respective group mean.

All participants initially deemed eligible were invited to the Douglas Research Centre for a behavioral testing session, where they completed a battery of standardized psychiatric and neuropsychological assessments. This included: the Mini-International Neuropsychiatric Interview (Sheehan et al., 1998), exclusion criteria = indications of undiagnosed psychiatric illness; the Beck Depression Inventory II (Beck et al., 1997), exclusion cut-off ≥19; and the Mini-Mental State Exam (Folstein et al., 1975), inclusion cut-off ≥26. After testing, participants donated blood samples for endocrine assessment and performed a practice version of the spatial context memory task in a mock MRI scanner. The mock session provided participants with an opportunity to familiarize themselves with the experimental setup (e.g., button responses). Only participants who met the inclusion/exclusion criteria and were able to perform the spatial context memory task in the mock MRI scanner were invited to participate in the second (MRI) session.

Upon arrival at the second session, female participants took a pregnancy test and were deemed eligible if the result was negative. Blood samples were again taken for endocrine assessment. In total, scanning lasted approximately 1.5-2 hours and included T1-weighted and T2-weighted structural MRI, diffusion MRI, resting-state fMRI, and task fMRI (during which the spatial context memory task was administered). The current study focuses only on the diffusion MRI and behavioral outputs of the task fMRI spatial context memory paradigm.

### 2.3 Endocrine assessments and menopause staging

We used STRAW+10 guidelines (Harlow et al., 2012) to categorize female participants as pre-, peri-, or post-menopausal based on their responses to a reproductive history/hormone use questionnaire, which was completed during the behavioral testing session and updated during the imaging session. Questions covered date of last period, number of periods in the past 12 months, and regularity of menstrual cycle, all of which form part of STRAW+10 criteria. Staging was then corroborated using estradiol-17β (E2) and follicle stimulating hormone (FSH) levels. These endocrine measures were assessed on plasma, derived from heparin blood collection tubes. Blood was drawn on non-fasting individuals by a certified research nurse during session two. Specimens were analyzed at the McGill University Health Centre (Glen site) Clinical Laboratories in Montreal. Endocrine Chemiluminescent immunoassays were performed on an Access Immunoassay System (Beckman Coulter) using the company’s reagents. Notably, for this sub-study, participants were not included if they were categorized as peri-menopausal or if their menopause status was indeterminate.

### 2.4 Spatial context memory paradigm

Participants completed a spatial context memory task during an fMRI scan (Fig. 1). The task was presented through E-Prime software (Psychology Software Tools, PA). Details of this task have been described elsewhere (Crestol et al., 2023). In brief, participants were asked to encode either six (easy spatial context memory task) or twelve (difficult spatial context memory task) face stimuli and their spatial location and to rate each face as either pleasant or neutral by pressing the corresponding button on the provided response boxes. The pleasantness rating was added to ensure participants deeply encoded the stimuli (Bernstein et al., 2002). Details regarding the timing and number of stimuli presented to participants are described in Fig. 1. After the encoding phase, participants received a break in which they rated how well they remembered the faces on a scale of 1 (very poorly) to 4 (very well). The break was 60 seconds long.

**Fig 1.**
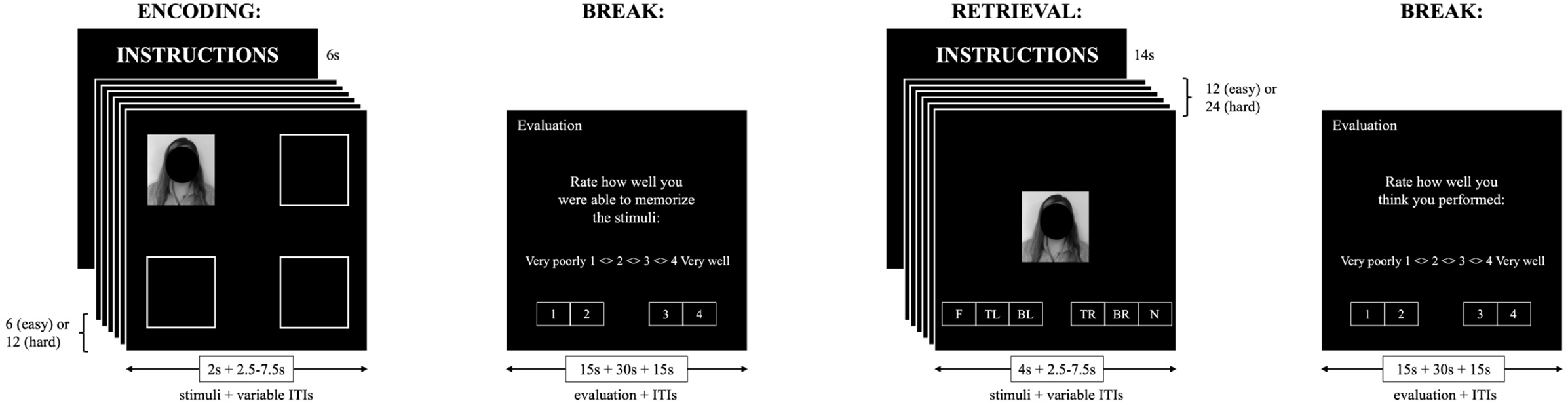
Spatial context memory paradigm. At encoding, participants were asked to encode face-location associations and make a pleasant/neutral decision for face stimuli presented in one of four quadrants. There was a one-minute break between encoding and retrieval. At retrieval, participants were presented with an equal number of previously encoded (“old”) and novel face stimuli in the center of the screen and asked to make a six-alternative forced-choice decision. Further details are presented in Section 2.4 (see also Crestol et al., 2023). Note that the face stimulus shown here is an illustrative example and was not part of the stimulus set. Moreover, it has been obstructed to comply with bioRxiv’s image policy. Abbreviations: ITIs = inter-trial intervals.

Following the break, participants performed the retrieval phase of the easy or hard task. During the easy task, participants were presented with the six previously encoded (“old”) face stimuli and six novel face stimuli, one at a time, in random order. During the hard task, participants were presented with twelve old and twelve novel face stimuli in random order. For each face, participants were instructed to respond by pressing one of six buttons corresponding to the following retrieval responses: (1) N – The face is NEW; (2) F – The face is FAMILIAR but I don’t remember its location;(3) TL – I remember the face and it was previously on the TOP LEFT; (4) BL – I remember the face and it was previously on the BOTTOM LEFT; (5) TR – I remember the face and it was previously on the TOP RIGHT; (6) BR – I remember the face and it was previously on the BOTTOM RIGHT. The mapping of buttons to response options was shown throughout. Participants were instructed not to guess and only respond (3) to (6) if they clearly recollected the face and its location. At the end of each retrieval phase, participants received a second break in which they were again asked to rate their performance. The break was 60 seconds long.

Participants could complete a maximum of four easy and four hard task runs, which were counterbalanced. Given that the easy task had half the number of face stimuli, this version was completed twice per run. In total, therefore, participants were exposed to a maximum of 96 faces at encoding (48 easy, 48 hard) and 192 faces at retrieval (96 easy [48 old, 48 novel], 96 hard [48 old, 48 novel]). However, due to participant withdrawal and software errors, not all participants experienced all runs. The minimum number of easy task runs completed was two, whereas the minimum number of hard task runs completed was three. As such, participants were exposed to a minimum of 70 faces at encoding (24 easy, 36 hard) and 120 faces at retrieval (48 easy [24 old, 24 novel], 72 hard [36 old, 36 novel]).

Participants were excluded from the current analyses if they provided zero correct spatial context retrieval responses or if they performed below chance in terms of correct rejections (1/6 = 0.16). Mirroring the approach taken by Crestol et al. (2023), we additionally ran regression analyses with age as a predictor of spatial context retrieval accuracy, item recognition, and misses on both the easy and hard task versions. Participants with a Cook’s D value more than three standard deviations from the group mean in three or more models were removed.

### 2.5 Diffusion MRI

#### 2.5.1 Acquisition and quality control

Scanning was conducted on a Siemens MAGNETOM 3T PrismaFit MRI scanner with a standard 32-channel head coil at the Douglas Research Centre. DTI analysis was conducted using diffusion MRI data acquired using the following sequence parameters: repetition time = 3100 ms; echo time = 105 ms; flip angle = 90°; phase-encoding direction = posterior-to-anterior; 63 slices; slice thickness = 2 mm; field of view = 130 x 130 mm; voxel dimensions = 2 x 2 x 2 mm; 65 volumes (64 *b* = 1000 s/mm^2^, 1 *b* = 0 s/mm^2^). Five non-diffusion-weighted volumes (*b* = 0 s/mm^2^) with reversed phase-encoding direction were also acquired for susceptibility distortion correction.

#### 2.5.2 Pre-processing

Diffusion MRI data were pre-processed in MRtrix3 (version 3.0.2; Tournier et al., 2019), incorporating tools from FSL (version 6.0.2; Jenkinson et al., 2012) and ANTs (version 2.3.1; http://stnava.github.io/ANTs/). Pre-processing followed the optimized diffusion pipeline described by Maximov et al. (2019). This included the following steps: noise correction; Gibbs ringing correction; EPI-induced geometric distortion correction; head motion, eddy current, and susceptibility distortion correction including the replacement of dropout slices; bias field correction; spatial smoothing with a 1 mm^3^ Gaussian kernel.

#### 2.5.3 Tract-based spatial statistics

Following pre-processing, diffusion tensor models were fit at each voxel using FSL’s DTIFIT. This produced parameter maps of FA and MD for each participant. Using these maps as input, the standard processing steps of tract-based spatial statistics (TBSS; Smith et al., 2006) were carried out. Briefly, participants’ FA maps were first aligned in common space (FMRIB58_FA template) using nonlinear registration. Next, a mean FA image was created and thinned (FA threshold = 0.2) to generate an FA skeleton. Each participant’s aligned FA data were then projected onto the FA skeleton. Finally, the TBSS non-FA pipeline was used to repeat these steps for MD, projecting MD data onto the FA skeleton.

#### 2.5.4 Region-of-interest segmentation

Binarized masks were generated for the 48 regions-of-interest (ROI) contained within the JHU ICBM-DTI-81 WM labels atlas. These binarized masks were then used to extract mean FA and MD values along the TBSS FA skeleton within each ROI, for each participant. Given limited head coverage in some participants, brainstem tracts were removed from consideration. Moreover, in the absence of a strong a priori hypothesis for hemispheric effects, we also opted to average values for left and right ROIs, where relevant. This left 21 WM ROIs for consideration (for more information, see Supplementary Material).

### 2.6 Statistical analyses

Behavioral analyses were conducted in R (version 4.1.3) using RStudio (version 2022.7.1.554). Brain-behavior analyses were conducted in MATLAB (version R2020) using open-source software (version 6.15.1; https://www.rotman-baycrest.on.ca/index.php?section=84).

#### 2.6. Behavioral analyses

Mean accuracy (rate correct, 0-1) and mean reaction time (RTs, ms) were calculated per participant for correct spatial context retrieval responses. Mean accuracy was calculated by summing the total number of correct face-location responses (i.e., options 3, 4, 5, and 6, see Fig. 1) and dividing by the total number of previously seen faces shown at retrieval. Mean RTs were calculated by summing the RTs for each correct face-location response and dividing by the total number of such responses. These measures were calculated separately for the easy and hard task and then standardized.

For our menopause analysis, we conducted multi-response partial least squares (PLS) regression analyses, implemented using the *plsreg2* function from the *plsdepot* package (version 0.2.0; Sanchez, 2023). This method handles multicollinearity among predictor variables, as is the case for menopause status and age, by creating a set of uncorrelated components to use in the model (Abdi, 2010). The aim of PLS regression is to predict Y (i.e., outcome variables) from X (i.e., predictor variables) and describe their common structure. To achieve this, X and Y are first decomposed to form a set of latent variables (LVs) that maximally explain the shared variance (i.e., cross-block covariance) between the original variables. The ability of a given LV to predict Y is then typically assessed via cross-validation, a process which produces Q^2^ values. LVs were considered significant and retained if Q^2^ ≥0.0975 (Abdi, 2010).

Although useful for dealing with multicollinearity among predictors, PLS regression does not provide insight as to whether age is differentially related to spatial context memory at specific stages of menopause. To address this, we conducted analyses using linear mixed-effects models (LMMs) for pre- and post-menopausal females, respectively. LMMs were fitted using the *lme4* package (version 1.1-29; Bates et al., 2015). Task difficulty (easy, hard) and age were entered as fixed effects, as was their interaction term. Task difficulty was coded using deviation coding. Age was centered and scaled. All LMMs included a random by-participant intercept. Statistical significance of fixed effects was determined via Satterthwaite approximations, implemented by the *lmerTest* package (version 3.1-3; Kuznetsova et al., 2017).

For our sex analysis, we sought to determine whether males and females differed in task performance, and to establish whether this was related to chronological age. Task difficulty (easy, hard), sex (male, female), and age were entered as fixed effects, as were the various interaction terms. For consistency, all other aspects of the LMMs were as described for menopause.

#### 2.6.2 Brain-behavior PLS analyses

For our brain-behavior analyses, we used a multivariate statistical technique: behavior PLS (bPLS; Krishnan et al., 2011; McIntosh & Lobaugh, 2004). Specifically, we used bPLS to examine whether chronological age was associated with WM microstructure within groups (i.e., pre-/post-menopause, male/female), before then examining if the resulting patterns were related to spatial context memory performance.

In each of these analyses, we first created two matrices: one containing the DTI data, the other containing the behavioral data. The DTI data matrix contained columns representing the 21 WM ROIs and rows representing standardized values for each participant nested within condition (i.e., FA, MD) and conditions nested within group. The behavioral data matrix contained standardized values of chronological age, with rows duplicated to mirror the nested structure of the DTI data matrix. These matrices were then cross-correlated and the output submitted to singular value decomposition, generating a set of LVs. For each LV, bPLS outputs a singular value, a correlation profile, and a singular image. The singular value represents the proportion of the cross-block covariance accounted for by each LV. The correlation profile (shown as a bar plot) highlights the association between “brain scores”, which signify the degree to which a given participant expressed the LV for each of the conditions, and the corresponding behavioral measure of interest. The singular image consists of positive and/or negative “saliences”, which identify whether ROIs are positively or negatively related to the correlation profile. The assignment of saliences as positive or negative is arbitrary.

Statistical significance of LVs was determined using permutation tests (2000 permutations). Specifically, for each LV, the probability that the permuted singular values exceeded the observed singular value was used to determine statistical significance (*p* < .05; McIntosh & Lobaugh, 2004). No correction for multiple comparisons was required as bPLS analyses are performed in one analytical step. The stability of saliences contributing to a given LV was assessed by dividing the observed value by its standard error (generated via 1000 bootstrap samples). The resulting ratio, referred to as the bootstrap ratio, is equivalent to a *z*-score and thus a higher value represents a more stable salience. In this study, a bootstrap ratio greater than ± 2.58 (corresponding to *p* < .01) was used to determine which ROIs reliably contributed to significant LVs.

For significant LVs, post-hoc regression analyses were performed on brain scores for conditions deemed to contribute to the identified pattern. These analyses were conducted to determine whether the expression of a given LV was related to the observed behavioral findings for both menopause status and sex. We first examined whether these brain scores were related to spatial context memory performance across all participants entered in a particular analysis. Thereafter, where relevant, we examined the association between brain scores and performance within groups.

### 2.7 Data and code availability

Code used for pre-processing and analysis is made publicly available via our Lab GitHub page (https://github.com/RajahLab/BHAMM_DWI_scripts/). Code reproducibility was assessed internally by multiple authors. Readers seeking access to data should email the Principal Investigator of the BHAMM Study, Professor Maria Natasha Rajah, for information.

## 3 Results

A total of 21 participants were excluded from our analyses. Reasons were use of hormonal birth control in post-menopause (*n* = 3), BMI > 40 (*n* = 1), previous hysterectomy and unilateral oophorectomy (*n* = 1), failed diffusion MRI quality control (*n* = 9), within-group age outlier (*n* = 2), and no correct spatial context retrieval responses (*n* = 5). Two male participants exhibited moderate-to-high levels of some cholesterol measures and were unmedicated but were nevertheless included due to their overall good health (determined via self-report) and satisfactory levels of neuropsychological and task performance. After applying these exclusions, therefore, the final sample consisted of 96 middle-aged adults (30 males, 32 pre-menopausal females, 34 post-menopausal females).

Demographic data separated by menopause status and sex are shown in Table 1. We used Welch’s *t*-tests and chi-square tests to examine group differences in continuous and categorical demographic variables, respectively. For pre-/post-menopause, we also examined differences in E2 and FSH. Starting with menopause status, we observed significant differences in age (*t*(61.126) = -17.415, *p* < .001), E2 (*t*(28.540) = 6.867, *p* < .001), and FSH (*t*(33.717) = - 16.392, *p* < .001). Hormonal differences were in the expected direction, such that pre-menopausal females had higher levels of E2 and lower levels of FSH. Age differences reflected the younger age of pre-menopausal females compared to post-menopausal females. This was expected given the interconnected nature of chronological and reproductive aging. For sex, we observed no statistically significant differences between males and females, although differences in age were near-threshold (*t*(69.908) = 1.947, *p* = .056).

**Table 1.** Demographic Characteristics Separated by Menopause Status and Sex.

|  | Pre-menopause (*n* = 32) | Post-menopause (*n* = 34) | Males (*n* = 30) | Females (*n* = 66) |
| --- | --- | --- | --- | --- |
| Age (years), M (SD) | 44.34 (2.66)*** | 57.70 (3.53)*** | 53.96 (5.87) | 51.22 (7.41) |
| Education (years), M (SD) | 16.44 (2.26) | 15.38 (2.65) | 16.73 (2.56) | 15.89 (2.51) |
| BMI (kg/m<sup>2</sup>), M (SD) | 24.80 (4.37) | 25.31 (3.95) | 25.02 (3.45) | 25.06 (4.13) |
| BDI-II, M (SD) | 4.19 (4.12) | 5.56 (5.51) | 4.97 (4.41) | 4.89 (4.90) |
| Handedness, % right | 90.62 | 91.18 | 80.00 | 90.91 |
| Antidepressants, *n* | 6 | 5 | 2 | 11 |
| Hormonal birth control, *n*<sup>a</sup> | 4 | 0 | NA | 4 |
| Polycystic ovarian syndrome, *n*<sup>a</sup> | 5 | 2 | NA | 7 |
| E2 (pmol/L), M (SD)<sup>b</sup> | 359.93 (215.42)*** | 76.17 (17.49)*** | - | 309.85 (223.76) |
| FSH (IU/L), M (SD)<sup>b</sup> | 7.70 (4.17)*** | 87.56 (27.56)*** | - | 50.90 (44.99) |

<sup>a</sup>Data were unavailable for 2 participants (1 pre-menopausal females, 1 post-menopausal females).
<sup>b</sup>Due to the detection thresholds of the ELISA used, E2 was measurable in 34 females (28 pre-menopausal females, 6 post-menopausal females) out of 61 females (28 pre-menopausal females, 33 post-menopausal females) who donated blood for hormonal analysis. FSH is given as average values for the 61 participants. While E2/FSH values were available for males, we do not report them as these measures were only used for menopausal staging (indicated by a dash [-]).
\**p* < .05, \*\**p* < .01, \*\*\**p* < .001.

### 3.1 Behavioral analyses

#### 3.1.1 Menopause

The PLS regression analysis identified one LV with a Q^2^ value above threshold (Q^2^ = 0.194). This LV accounted for 95.36% of the variance in X (i.e., menopause status, age) and 21.68% of the variance in Y (i.e., mean accuracy, mean RTs). Fig. 2A depicts the correlations between the original variables and the LV. Menopause status and age were both negatively associated with the LV, as were mean RTs on the easy and hard versions of the task. By contrast, mean accuracy for the easy and hard versions of the task both correlated positively with the LV. Together, these results suggest that the LV captured a pattern in which age and post-menopause status were both positively correlated with mean RTs but negatively correlated with mean accuracy.

**Fig 2.**
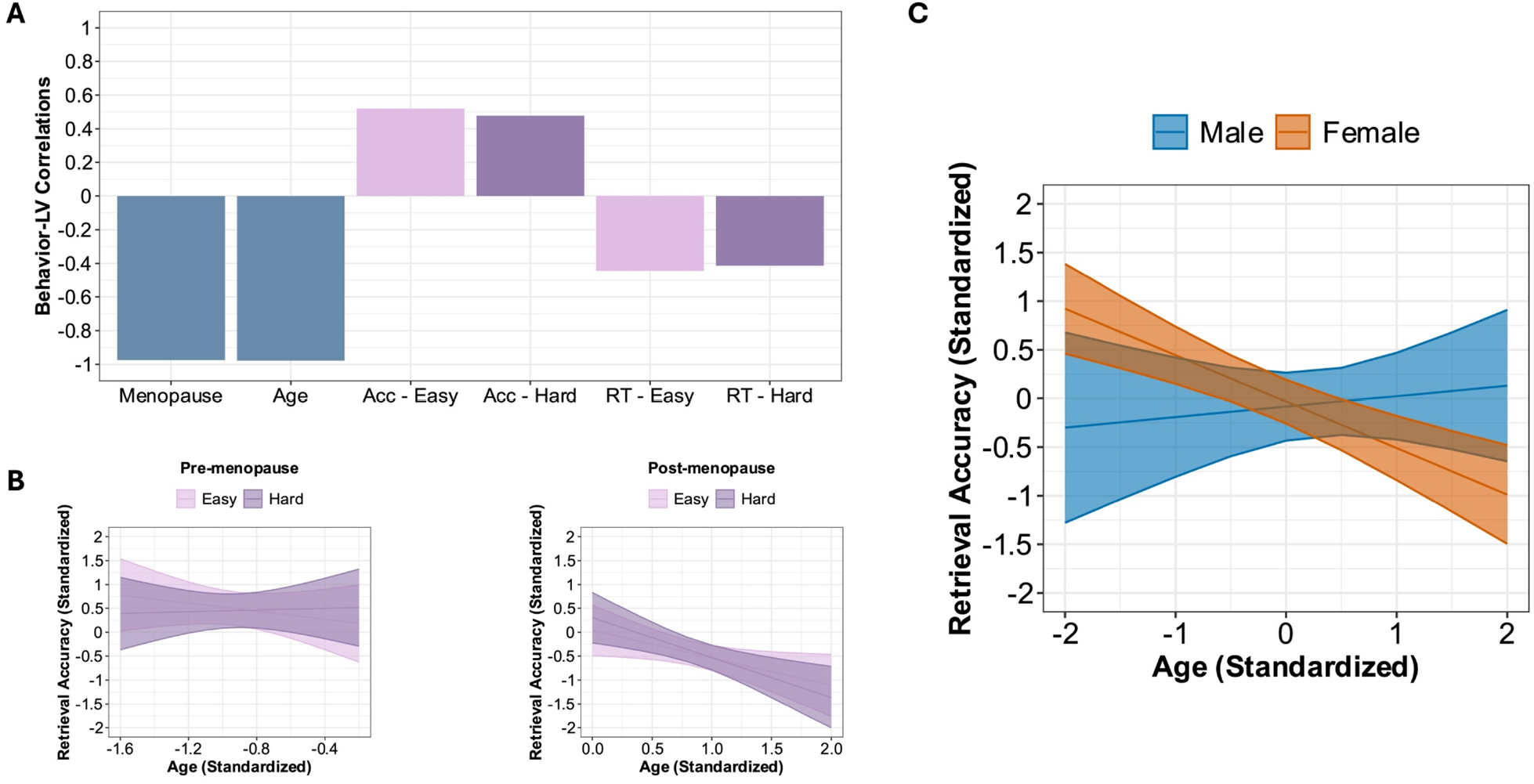
Menopause status- and sex-related differences in age associations with spatial context retrieval accuracy. **A)** Correlation profile for the LV identified by PLS regression. The bars reflect the correlation between the original independent (blue)/dependent (pink/purple) variables and the LV. **B)** Association between age (standardized) and spatial context retrieval accuracy (standardized) as a function of task difficulty (easy, hard) in pre-menopausal females (left) and post-menopausal females (right). Lines of best fit and 95% confidence intervals are based on predicted values. **C)** Association between age (standardized) and spatial context retrieval accuracy (standardized) in males and females. Lines of best fit and 95% confidence intervals are based on predicted values. Abbreviations: Acc = spatial context retrieval accuracy, RT = reaction times (correct spatial context retrieval responses), LV = latent variable.

We subsequently conducted separate LMMs for mean accuracy within groups (Fig. 2B). Among pre-menopausal females, there were no statistically significant associations between mean accuracy and task difficulty (*F*(1, 30) = 1.377, *p* = .250), age (*F*(1, 30) = 0.133, *p* = .718), or their interaction (*F*(1, 30) = 1.904, *p* = .178). Among post-menopausal females, mean accuracy was significantly associated with age (*F*(1, 32) = 8.373, *p* = .007) but not task difficulty (*F*(1, 32) = 1.563, *p* = .220) or their interaction (*F*(1, 32) = 1.450, *p* = .237). The age-accuracy association was negative, reflecting lower accuracy with advanced age (*β* = -0.706). Thus, while the PLS regression analysis indicated that age and post-menopausal status were both negatively related to spatial context retrieval accuracy, further analyses showed that age associations varied by menopause status. Specifically, age negatively correlated with accuracy in post-but not pre-menopausal females. Mirroring the accuracy analysis, separate LMMs were conducted for mean RTs in each menopausal group. Among pre-menopausal females, there were no significant associations between mean RTs and task difficulty (*F*(1, 30) = 0.164, *p* = .688), age (*F*(1, 30) = 0.775, *p* = .386), or their interaction (*F*(1, 30) = 0.262, *p* = .613). Among post-menopausal females, the same overall pattern was evident: mean RTs were not significantly associated with task difficulty (*F*(1, 32) = 0.315, *p* = .579), age (*F*(1, 32) = 0.375, *p* = .545), or their interaction (*F*(1, 32) = 0.523, *p* = .475).

#### 3.1.2 Sex

For sex, we found that mean accuracy was not significantly related to task difficulty (*F*(1, 92) = 0.001, *p* = .978), age (*F*(1, 92) = 2.220, *p* = .140) or sex (*F*(1, 92) = 0.039, *p* = .843).

However, there was a significant interaction between age and sex (*F*(1, 92) = 8.348, *p* = .005). This interaction (Fig. 2C) reflected a negative association between age and accuracy in females (*β* = -0.475, *p* < .001) but not males (*β* = 0.152, *p* = .431). No other interactions were statistically significant (all *p* ≥.584). For mean RTs, there were also no significant associations with task difficulty (*F*(1, 92) = 0.556, *p* = .458), age (*F*(1, 92) = 2.534, *p* = .115) or sex (*F*(1, 92) = 0.363, *p* = .549). Consistent with the mean accuracy analysis, we observed a significant interaction between age and sex for mean RTs (*F*(1, 92) = 4.614, *p* = .034). This interaction reflected a positive association between age and mean RTs in females (*β* = 0.419, *p* < .001) but not males (*β* = -0.062, *p* = .754). No other interactions were statistically significant (all *p* ≥.097).

Given the notable difference in sample size and near-significant difference in age between groups, we repeated our analyses using a subset of females that were matched on age (and education) to the available males. These analyses were intended to assess the sensitivity of our results. Overall, our results remained unchanged (see Supplementary Material). Considering these findings, we opted to conduct our brain-behavior PLS analyses on the full sample.

### 3.2 Behavior PLS analyses

#### 3.2.1 Menopause

The bPLS analysis examining the association between age and WM microstructure as a function of menopause status identified one statistically significant LV (LV1_meno_; *p* = .036, 62.34% cross-block covariance explained). LV1_meno_ identified WM tracts in which advanced age was associated with lower FA and higher MD in both pre- and post-menopausal females (Fig. 3A). The anterior corona radiata, body of the corpus callosum, and genu of the corpus callosum contributed reliably to LV1_meno_ (Fig. 3B).

**Fig 3.**
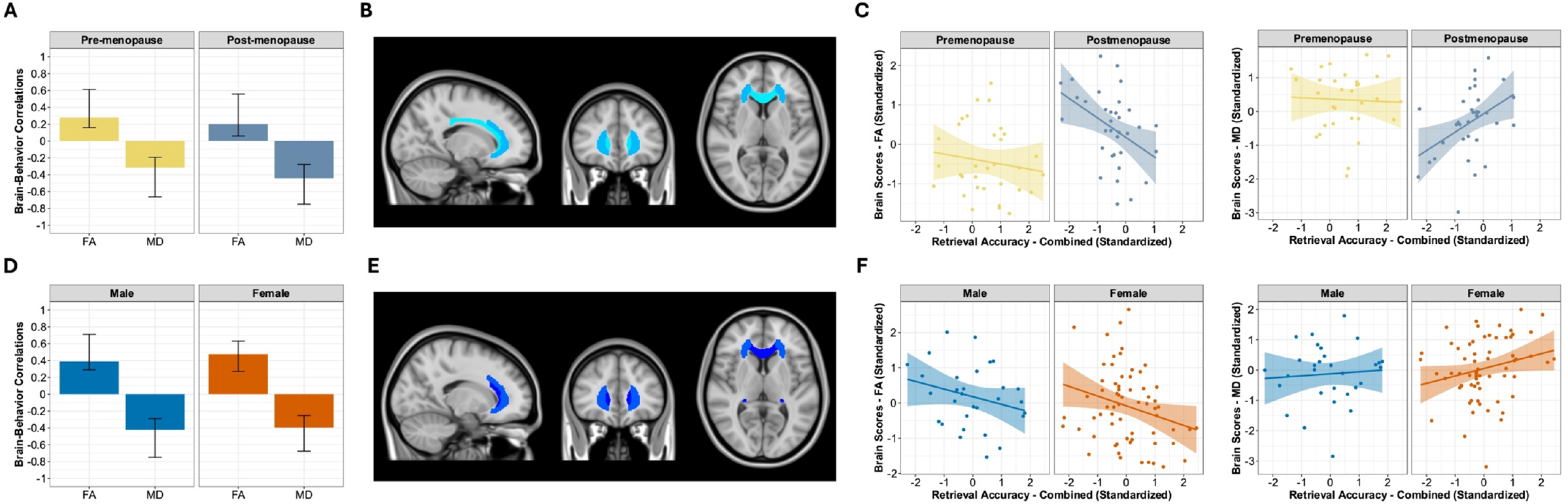
Menopause status- and sex-related differences in age associations with WM and WM-spatial context retrieval accuracy. **A)** Brain-behavior correlation profile for LV1_meno_. The bars reflect the correlation between age and FA/MD in pre- and post-menopausal females. The whiskers on the bars reflect 95% confidence intervals based on bootstrap results. **B)** Singular image for LV1_meno_. Only negative salience regions were identified and are colored in blue (darker = stronger BSR value). These negative saliences identify WM tracts that were negatively associated with the correlation profile. **C)** Correlation plots showing the association between brain scores (standardized) for LV1_meno_ (FA, MD) and spatial context retrieval accuracy collapsed across easy and hard task versions (standardized). **D)** Brain-behavior correlation profile for LV1_sex_. The bars reflect the correlation between age and FA/MD in males and females. The whiskers on the bars reflect 95% confidence intervals based on bootstrap results. **E)** Singular image for LV1_sex_. Only negative salience regions were identified and are colored in blue (darker = stronger BSR value). These negative saliences identify WM tracts that were negatively associated with the correlation profile. **F)** Correlation plots showing the association between brain scores (standardized) for LV1_sex_ (FA, MD) and spatial context retrieval accuracy collapsed across easy and hard task versions (standardized). Abbreviations: BSR = bootstrap ratio, FA = fractional anisotropy, MD = mean diffusivity, LV = latent variable, WM = white matter.

Given that age was associated with FA and MD in both groups, we examined associations between LV1_meno_ brain scores for both conditions and mean accuracy averaged across easy and hard task versions. We focused on this combined measure of performance due to a lack of task difficulty effects. Regression analyses revealed that LV1_meno_ brain scores for FA (*β* = -0.422, *p* < .001) and MD (*β* = 0.321, *p* = .009) were both significantly related to mean accuracy averaged across task versions. Interestingly, when analyzed within group, we found that this pattern was evident among post-menopausal females (FA: *β* = -0.352, *p* = .014; MD: *β* = 0.342, *p* = .011) but not pre-menopausal females (FA: *β* = - 0.151, *p* = .449; MD: *β* = -0.046, *p* = .818) (Fig. 3C). Thus, given that the saliences associated with LV1_meno_ were negative, these analyses provide evidence that the expression of this LV (lower FA and higher MD with advanced age) was related to lower spatial context retrieval accuracy in post-menopausal females.

#### 3.2.2 Sex

The bPLS analysis examining the association between age and WM microstructure as a function of sex identified one statistically significant LV (LV1_sex_; *p* < .001, 62.68% cross-block covariance explained) and one near-significant LV (LV2_sex_; *p* = .052, 23.97% cross-block covariance explained; for more information, see Supplementary Material). LV1_sex_ identified WM tracts in which advanced age was associated with lower FA and higher MD in both males and females (Fig. 3D). The WM tracts that reliably contributed to this pattern were the anterior corona radiata, fornix/stria terminalis, and genu of the corpus callosum (Fig. 3E).

As in the menopause status analysis, we examined associations between LV1_sex_ brain scores and mean accuracy averaged across task versions. Regression analyses revealed that LV1_sex_ brain scores for FA (*β* = -0.26, *p* = .011) but not MD (*β* = 0.185, *p* = .071) were significantly related to mean accuracy. Subsequent within-group analyses indicated that this was evident in females (*β* = -0.240, *p* = .036) but not males (*β* = -0.323, *p* = .157), although the observed patterns were qualitatively similar, perhaps due to the inclusion of pre-menopausal females (Fig. 3F). As the saliences associated with this LV were negative, these analyses suggest that the expression of this pattern (lower FA with advanced age) was related to lower spatial context retrieval accuracy, and that – statistically – this was identified in females but not males..

## 4 Discussion

The current study investigated menopause status- and sex-related differences in the association between age, spatial context memory, and white matter microstructure at midlife. Consistent with recent work (Crestol et al., 2023), advanced age was associated with lower spatial context retrieval accuracy (independent of task difficulty) among post-but not pre-menopausal females. To further assess whether this association was driven by menopause and not early-vs. late-midlife effects, we then examined whether females – irrespective of menopause status – exhibited a different pattern to males. In contrast to prior lifespan studies (Subramaniapillai et al., 2019, 2022; Wang et al., 2022), we found that advanced age was indeed associated with lower spatial context retrieval accuracy in females but not males. To our knowledge, this is the first study to identify sex differences in the relationship between age and episodic memory, assessed using item-location spatial context memory paradigms, in middle-aged adults. Taken together, our results suggest that the transition from pre-to post-menopause is associated with an increased vulnerability to age-related changes in the brain and cognition, and that this vulnerability appears to contribute to sex differences in age-related episodic memory decline at midlife.

Declining ovarian hormone production during the menopausal transition offers a plausible explanation for these results. The transition from pre-to post-menopause is characterized by significant hormonal changes, including a marked decrease in estrogen levels, most notably E2 (Harlow et al., 2012). More than two decades of research in rodents and non-human primates has demonstrated that E2 acts on numerous brain regions, including the hippocampus and prefrontal cortex, influencing memory function (Jacobs & Goldstein, 2018). Emerging reports in humans complement this work, leading some to propose that menopause-related reductions in E2 increase vulnerability to age-related cognitive decline and dementia risk (e.g., Rahman et al., 2019). While we were unable to examine the influence of E2 directly in this study, owing to the detection threshold of the ELISA used, our findings are nevertheless consistent with this proposal.

At the neural level, advanced age was associated with lower FA and higher MD, particularly among the anterior corona radiata and genu of the corpus callosum, independent of menopause status and sex. These findings imply that age-related decline in WM microstructure is evident at midlife, notably among regions of frontal WM. This is consistent with an anterior-posterior gradient of age-related decline in WM microstructure, whereby anterior (i.e., frontal) tracts exhibit greater vulnerability to age effects (Madden & Parks, 2016). However, the lack of menopause- and sex-related differences was inconsistent with our original hypothesis. One possible explanation is that we did not have sufficient statistical power to detect these effects, owing to our modest sample size. That said, a recent study of 812 middle-aged adults also failed to observe sex differences in the effect of age on DTI measures (Eikenes et al., 2022). It is possible, therefore, that sex differences in age-WM associations are detectable only when including participants from a broader range of the adult lifespan (e.g., Isaac Tseng et al., 2021). Such an explanation does not account for the failure to detect menopause-related differences in age-WM associations, although this is the first study to directly address this question. Only one other study to our knowledge has investigated menopause status effects on WM (Mosconi et al., 2021) and these authors opted to include age as a covariate. As such, the lack of menopause-related differences in the association between age and WM microstructure is not necessarily incongruent with their results.

Intriguingly, while advanced age was associated with microstructural variability in the WM of both males and females, subsequent analyses showed that the expression of this pattern was related to spatial context retrieval accuracy in post-menopausal females. That is, among post-menopausal females, lower FA/higher MD with advanced age was associated with poorer spatial context retrieval accuracy. These findings add weight to the view that the transition from pre-to post-menopause may increase vulnerability to the effects of age on the brain and episodic memory at midlife (Crestol et al., 2023). Furthermore, given that decline was largely restricted to frontal WM, it is possible that this vulnerability may contribute to increased age-related activation in prefrontal and inferior parietal cortices among females (Subramaniapillai et al., 2019), potentially serving as a compensatory mechanism.

The current study has limitations that should be considered when interpreting our results. First, our pre-menopausal females were not scanned during a specific phase of their menstrual cycle, as done in some related studies (e.g., Jacobs et al., 2016). It is possible, therefore, that menstrual cycle-related variability in the structure and/or function of regions involved in episodic memory (Dubol et al., 2021; Zsido et al., 2023) may have contributed to the lack of age effects among this group. However, recent research showing that verbal and spatial cognition remain relatively stable across the menstrual cycle casts some doubt on this, instead suggesting that such variability – if present – is adaptive and may not necessarily influence cognitive function (Pletzer et al., 2024). Second, while it is our view that chronological and reproductive aging constitute synergistic processes, we were unable to comprehensively examine age-independent effects of menopause status due to the observed age differences between our pre- and post-menopausal groups. Future research would benefit from the inclusion of larger samples with greater overlap in age between menopausal groups, facilitating the examination of both synergistic and independent effects. Third, we did not examine the role of white matter hyper-intensities (WMHs) in our sample. WMHs are common in midlife (d’Arbeloff et al., 2019) and overall WMH burden accelerates post-menopause (Lohner et al., 2022). It is possible, therefore, that group differences in WMH burden may have been evident. As prior reports indicate that WMHs impact DTI-derived measures (Svärd et al., 2017) and are more strongly associated with episodic memory than FA (Lockhart et al., 2012), we cannot rule out the possibility that our findings were influenced by their presence. Fourth, as is the case for other neuroimaging studies of menopause (e.g., Crestol et al., 2023; Jacobs et al., 2016), this study was cross-sectional. While this approach has produced interesting insights, longitudinal studies are needed to refine our understanding of the ways in which the menopausal transition influences the brain and episodic memory.

## 5 Conclusions

In this study, we found that the relationship between chronological age and spatial context memory differed as a function of menopause status and sex. Advanced age was associated with lower retrieval accuracy in post-but not pre-menopausal females and in females (collapsed across menopause status) but not males. We additionally observed that the degree to which post-menopausal females exhibited age-related decline in frontal WM was linked with lower spatial context retrieval accuracy. These results suggest that the menopausal transition – a major midlife event in females – increases vulnerability to age effects on the brain’s WM and episodic memory. This study thus adds to a growing body of literature highlighting how menopause may act as a critical period in female brain aging and underscores the importance of midlife in neurocognitive aging.

## Acknowledgements

We would like to thank the research participants of the Brain Health at Midlife and Menopause (BHAMM) study for their time and contribution to science. We acknowledge the support of part-time research assistants (H. Azizi, R. Young, A. Condescu, L. Khyatt, M. Zhu), and trainees (G. Velez Largo, A. Duval, J. Snytte, S. Loparco) who assisted with participant recruitment, testing, or MRI quality control. We are also thankful for the support of our BHAMM study collaborators (E. Jacobs, G. Einstein, R. K. Olsen, P. Rosa-Neto, D. Titone); Team 9, 10, and WSGD Theme of the Canadian Consortium on Neurodegeneration in Aging (CCNA); the MRI staff at the CIC, Douglas Research Centre; and Dr. D. Cohen for help with recruitment.

## CRediT author statement

Rikki Lissaman: Conceptualization; Data curation; Formal analysis; Methodology; Software; Validation; Visualization; Writing – Original draft; Writing – Review & editing. Sricharana Rajagopal: Data curation; Formal analysis; Methodology; Software; Validation; Visualization. Julia Kearley: Investigation; Software; Validation; Writing – Review & editing. Stamtoula Pasvanis: Data curation; Investigation; Project administration; Methodology; Validation. Maria Natasha Rajah: Conceptualization; Formal analysis; Funding acquisition; Methodology; Project administration; Resources; Supervision; Writing – Original draft; Writing – Review & editing.

