## Supplementary Material for "Menopause status- and sex-related differences in age associations with spatial context memory and white matter microstructure at midlife"

### **S1. White Matter Regions-of-Interest**

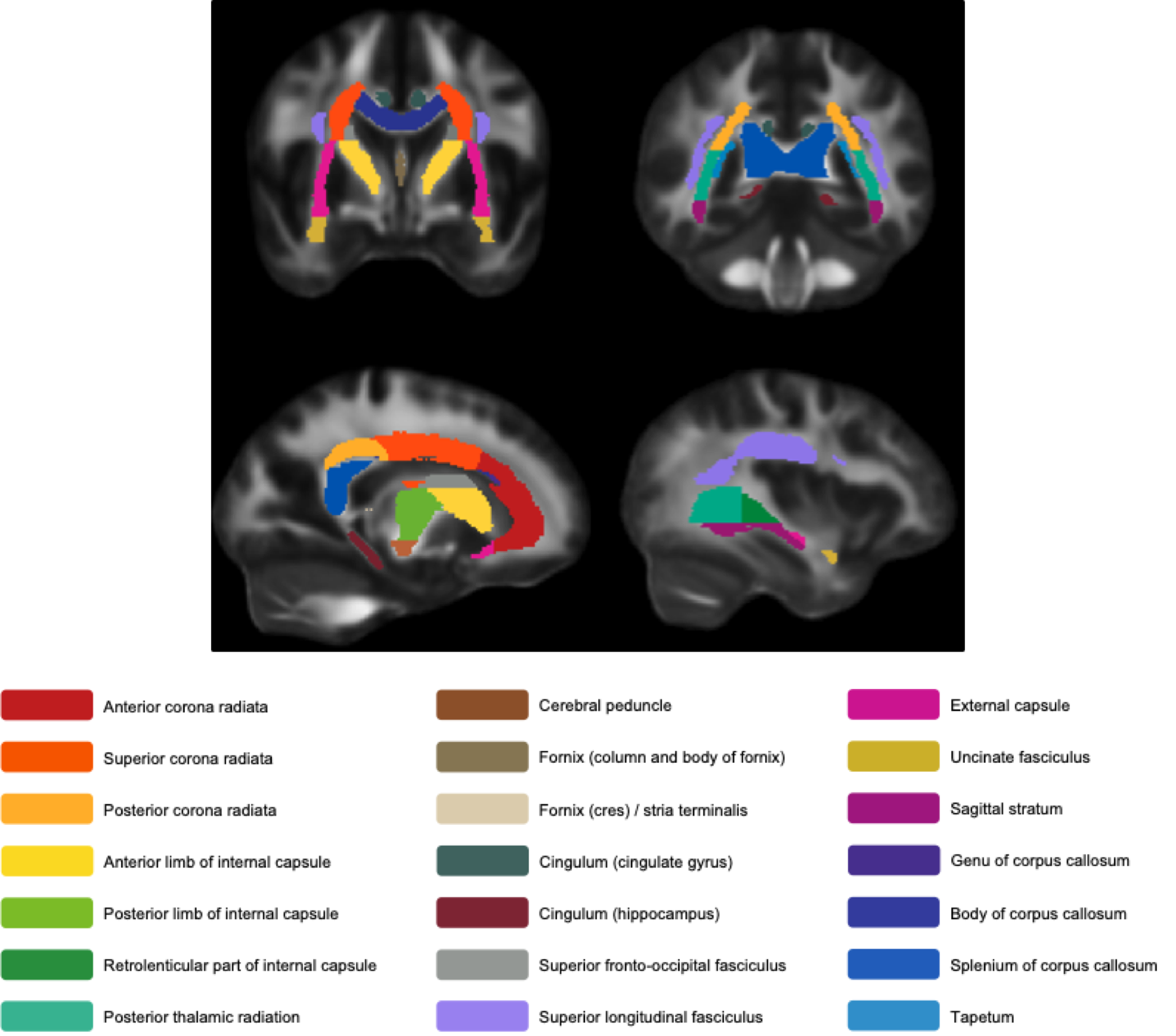

*Note*. Locations of the 21 ROIs from the JHU ICBM-DTI-81 WM labels atlas (Mori et al., 2008; Oishi et al., 2008). All ROIs are shown bilaterally, as we averaged values across hemispheres (where applicable).

### **S2. Age- and Education-Matched Sex Analysis**

Our female group included more participants than our male group (*n* = 66 vs. *n* = 30) and covered a wider age range (40.07-65.14 years vs. 43.84-65.46 years). As such, we decided to repeat our sex-related behavioral analyses using a new, smaller sample of females (*n* = 30; 8 pre-menopausal, 22 post-menopausal) that were matched in age and education to the available males. The goal of the analysis was to check the sensitivity of our results to these differences in age characteristics. We additionally opted to match on education due to concerns that cognitive reserve may also be relevant.

Matching was conducted in R using optimal pair matching (a.k.a., optimal matching), implemented via the MatchIt package (version 4.4.0; Ho et al., 2011). Age and education data for the male and *new* female group are shown in the Supplementary Table below. There were no statistically significant differences between groups in age (*t*(54.492) = 0.181, *p* = .857) or education (*t*(57.964) = -0.613, *p* = .542). The standardized effect sizes, shown in the Supplementary Table, were also small. However, as frequentist null hypothesis statistical tests (NHSTs) cannot provide evidence in favour of the null (in this case, no difference between groups), we additionally calculated Bayes Factors (BFs) in R using the BayesFactor package (version 0.9.12-4.7; Morey & Rouder, 2024). BFs enable researchers to assess the degree to which observed data favours one of two hypotheses – e.g., the null (H_0_) and the alternative (H_1_) (for an accessible introduction, see Quintana & Williams, 2018). According to Lee and Wagenmakers (2014), BFs greater than 3 provide at least moderate evidence for the alternative hypothesis, while BFs less than 0.33 provide at least moderate evidence for the null hypothesis. In our matched sample, we observed moderate evidence in support of the null for age (BF_10_ = 0.266 [0.01% error]) and anecdotal-to-moderate evidence in support of the null for education (BF_10_ = 0.30 [0.01% error]). Overall, therefore, the matching procedure appeared to have been successful.

| **Supplementary Table**  *Demographic Characteristics for Males and Age-/Education-Matched Females* | | | |
| --- | --- | --- | --- |
|  | Males (*n* = 30) | Females (*n* = 30) | Effect size (Cohen’s *d*) |
| Age (years)  *M* (*SD*)  Range | 53.96 (5.87)  43.84-65.46 | 54.28 (7.61)  40.63-65.14 | 0.047  NA |
| Education (years)  *M* (*SD*)  Range | 16.73 (2.56)  11-20 | 16.33 (2.50)  13-20 | 0.158  NA |

To determine whether our results were robust, we then repeated our LMMs. All aspects mirrored those reported in the manuscript. For mean accuracy, there were no significant associations with task difficulty (*F*(1, 56) < .001, *p* = .995), age (*F*(1, 56) = 1.136, *p* = .291), or sex (*F*(1, 56) = 0.176, *p* = .677). Nevertheless, we did observe a significant interaction between age and sex (*F*(1, 56) = 5.074, *p* = .028). This interaction reflected a negative association between age and accuracy in females (*β* = -0.407, *p* = .009) but not males (*β* = 0.146, *p* = .457). No other interactions were statistically significant (all *p* ≥ .427). For mean RTs, we again found no statistically significant associations with task difficulty (*F*(1, 56) < .001, *p* = .998), age (*F*(1, 56) = 2.784, *p* = .101), or sex (*F*(1, 56) = 0.054, *p* = .817). Mirroring the mean accuracy analysis, however, we did observe a significant interaction between age and sex for mean RTs (*F*(1, 56) = 4.536, *p* = .038). This interaction reflected a positive association between age and mean RTs in females (*β* = 0.460, *p* = .003) but not males (*β* = -0.056, *p* = .772). No other interactions were statistically significant (all *p* ≥ .343).

In sum, there were no notable differences between these analyses and those reported in the manuscript. It appears unlikely, therefore, that our behavioral results for sex were appreciably influenced by differences in sample size, age range, or years of education.

**S3. BPLS: Sex Analysis**

Our sex-based bPLS analysis identified a second, near-significant LV (LV2_sex_; *p* = .052, 23.97% cross-block covariance explained). LV2_sex_ identified WM tracts in which there was a DTI parameter-by-group interaction in the effect of age (shown below). More specifically, this LV identified tracts in which age was negatively associated with FA (but not MD) in females and negatively associated with MD (but not FA) in males. Five WM tracts reliably contributed to this pattern: the cingulum (hippocampus), cerebral peduncle, external capsule, fornix, and posterior limb of the internal capsule.

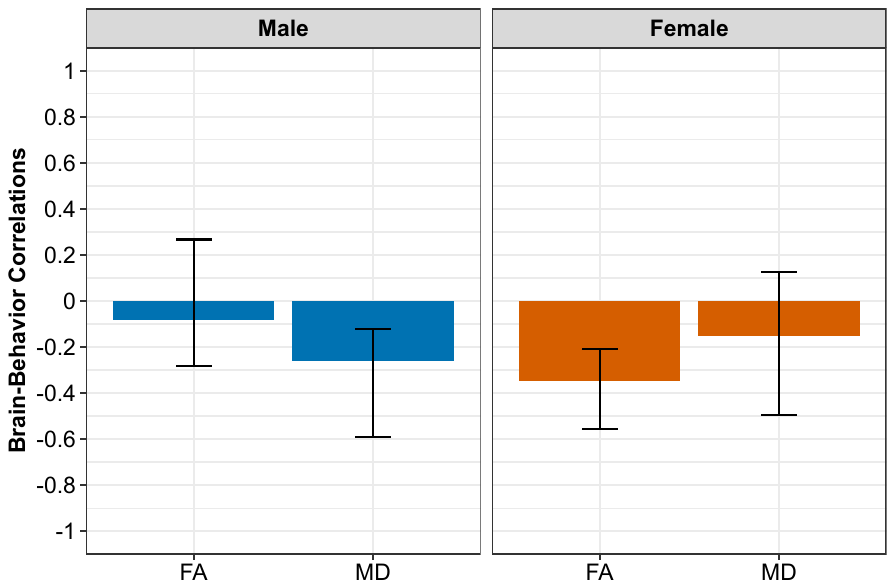

*Note*. Correlation profile for LV2_sex_.
